# Cytoplasmic folding, mis-folding, and early stages of aggregation

**DOI:** 10.1101/2022.10.23.513428

**Authors:** Premila P. Samuel Russell, Meredith M. Rickard, Mayank Boob, Martin Gruebele, Taras V. Pogorelov

**Affiliations:** Department of Chemistry, University of Illinois at Urbana-Champaign, Urbana, Illinois 61801, USA; Center for Biophysics and Quantitative Biology, University of Illinois at Urbana-Champaign, Urbana, Illinois 61801, USA; Beckman Institute for Advanced Science and Technology, University of Illinois at Urbana-Champaign, Urbana, Illinois 61801, USA; Department of Physics, University of Illinois at Urbana-Champaign, Urbana, Illinois 61801, USA; National Center for Supercomputing Applications, University of Illinois at Urbana-Champaign, Urbana, Illinois 61801, USA; School of Chemical Sciences, University of Illinois at Urbana-Champaign, Urbana, Illinois 61801, USA

## Abstract

We examine how cellular interactions in an all-atom model of the *Homo sapiens* cytoplasm influence the early folding events of Protein B (PB), a three-helix bundle protein. While PB is known to fold during *in vitro* simulations in three microseconds, all three initially unfolded PB copies in our cytoplasm model never completely reached their native topology across our 31 microsecond simulation. We were able to capture initial formation of all three helices and a compact topology similar to the native state. Sticking interactions between PB and surrounding macromolecules, as well as other unfolded PBs, became competitive with PB folding. Interaction between PB copies seeded beta-strand formation, modeling initial events of protein aggregation. Finally, the fold-switching potential of PB related GA domains has been explored in previous studies, and the sticking and crowding in our model thus initiates sampling of helix/sheet structural plasticity of PB.

---

Past *in vitro* folding studies have established that the information necessary for spontaneous protein (re)folding is stored in the primary sequence of protein domains, although folding yield may decrease due to frustration as protein size grows^1,2^. However, interaction between the complex heterogenous cellular environment and a protein can lead to chaperoned folding, misfolding and even fold switching^3–6^. To understand at an atomic level how a *Homo sapiens* cellular environment influences protein folding trajectories, we placed three unfolded copies of Protein B (PB), a fast-folding three-helix bundle, within an all-atom cytoplasm model of the human U-2 OS cell line (Fig. 1A). Our 47-residue monomeric PB protein construct originates from the bacterial sequence of *Peptostreptococcus magnus* protein B’s albumin binding GA domain^7^ (sequence in SI). During our 31-μs simulation^8^ of PB in the U-2 OS cytoplasm model, we were able to model early events of polypeptide folding, misfolding induced by protein-protein interaction, and seeding of β-strands in PB homo-oligomers, the earliest stage of intracellular protein aggregation.

**Fig 1.**
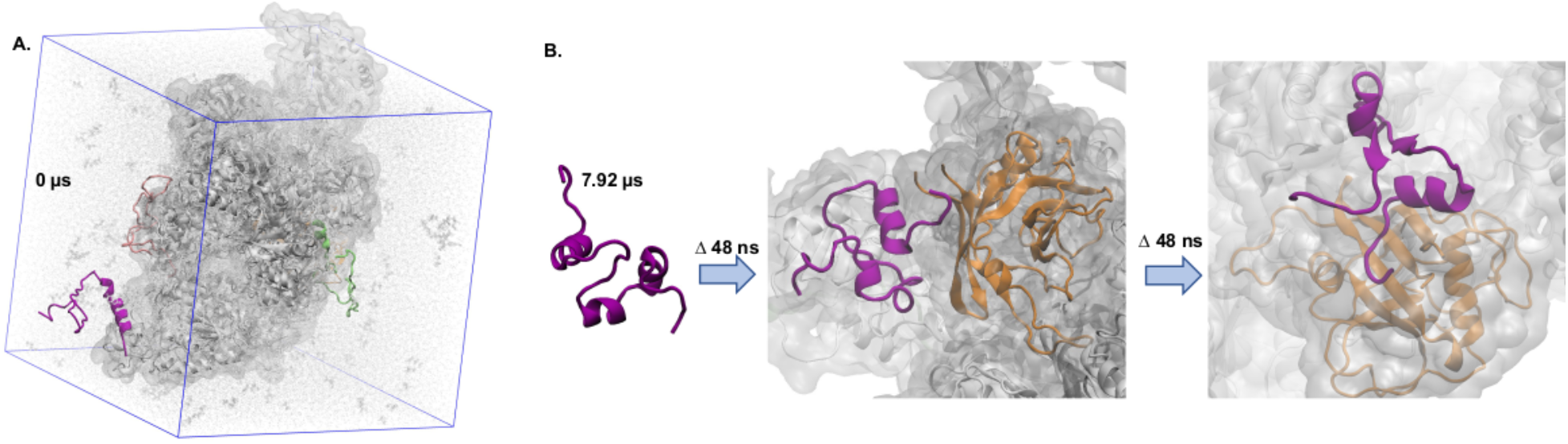
Snapshots of the transition between folding and misfolding events of PB in the cytoplasm model. **(A)** Three unfolded copies of PB after equilibration in our cytoplasm model. **(B)** A PB copy formed native-like structure, represented by three collapsed α-helical strands, and then started forming non-native β-strands following a sticking event between the partially unfolded N-terminus region of Helix1 and cyclophilin G. Protein backbones of the three individual PB copies are represented as a green (PB1), a pink (PB2), and a purple cartoon (PB3), highlighting changes in secondary structure. Volumetric isosurfaces of all other macromolecules are shown as transparent gray surface representations. Cyclophilin G is represented as an orange cartoon. All other macromolecular backbones are colored as grey cartoons. In (A), water molecules are shown as white transparent discs and metabolites are shown as gray transparent licorices. The clumping of macromolecules is indicative of over-stickiness of the force field, discussed previously.^10,11^.

In our previous tens of μs cellular simulation studies of unfolded WW domain, another canonical small fast-folding protein characterized by a triple-stranded β sheet fold, we observed the WW domain becoming trapped in a mis-folded state, driven by excessive protein-protein sticking in *E. coli* cytoplasm models^4^. In *in vitro* simulations on a comparable time scale, the same protein undergoes complete folding events in under 10 μs on average^9^.

In another *in vitro* study, PB was seen to fold to its native structure in 3 μs on average^12^. In our present mammalian cytoplasm model, we observe all three unfolded PB copies (PB1, PB2, and PB3) sampling a wide space of conformations (Figs. 2, 3, S1, S2), relative to the *in vitro* study of PB^12^. The less crowded U-2 OS cell cytoplasm at ~280 mg/ml macromolecular concentration^8^, relative to the ~300 mg/ml in the *E.coli* cell cytoplasm^4^, likely is the cause of the reduced lifetime of protein sticking events in our present U-2 OS cytoplasm, which we analyzed in our previous work^8^. Reduced protein-protein contact lifetimes allow for more configurational space exploration. The overall negative charge of the PB protein, *vs.* the WW domain’s positive charge, also contributed: the cytoplasmic macromolecules carry net negative charge^13^. Meanwhile, the study on the structural plasticity potential of GA domains by Orban and co-workers has demonstrated GA domain fold switching to immunoglobulin-binding GB folds of 4β + α, which can be induced by a single critical mutation^14^. In our current study we do observe long-lived β strand elements in both PB1 and PB3 from about 1-7 μs (Figs. 3A, 3C).

**Fig 2.**
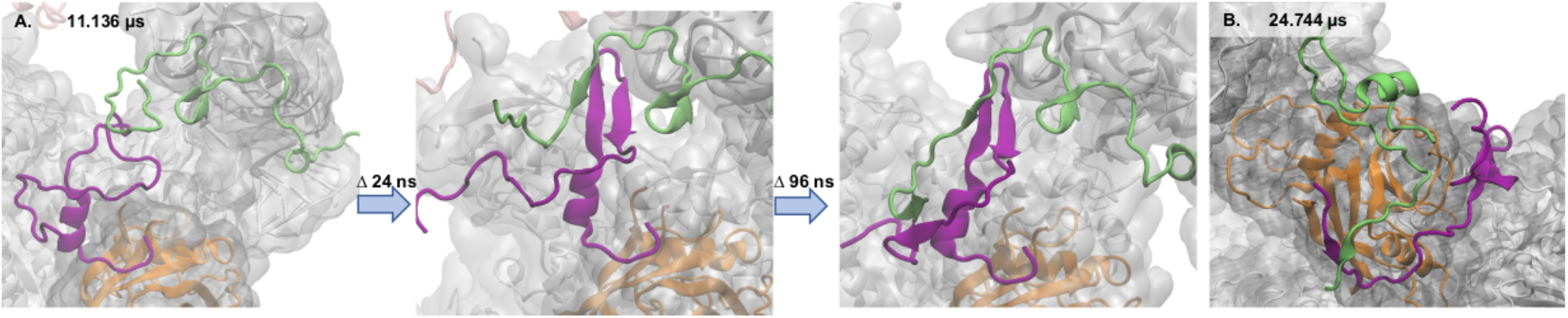
Sticking and aggregating. **(A)** Seeding of β-strands at interacting surfaces between two monomeric PB copies (PB1 and PB3). **(B)** Simultaneous sticking and aggregating events for protein B copies.

At about 7 μs into the simulation, all three helices form simultaneously in PB3 (Fig. 3C), and at 7.92 μs, PB3 approached the topology of a collapsed helix bundle (Figs. 1B, 3C, 3F and 4A), with a radius of gyration *(R_g_)* of 1.21 nm (close to folded PB’s *R_g_* of 1.03 nm). However, PB3 does not reach its native-like topology during our cellular cytoplasm simulation, with only 40% of native contacts formed (Fig. S2) and with a C-α root-mean-square-deviation (RMSD) of 1.04 nm (Fig. 4D). The three helices arrange in a compact, but not native, topology (Fig. 3F).

**Fig 3.**
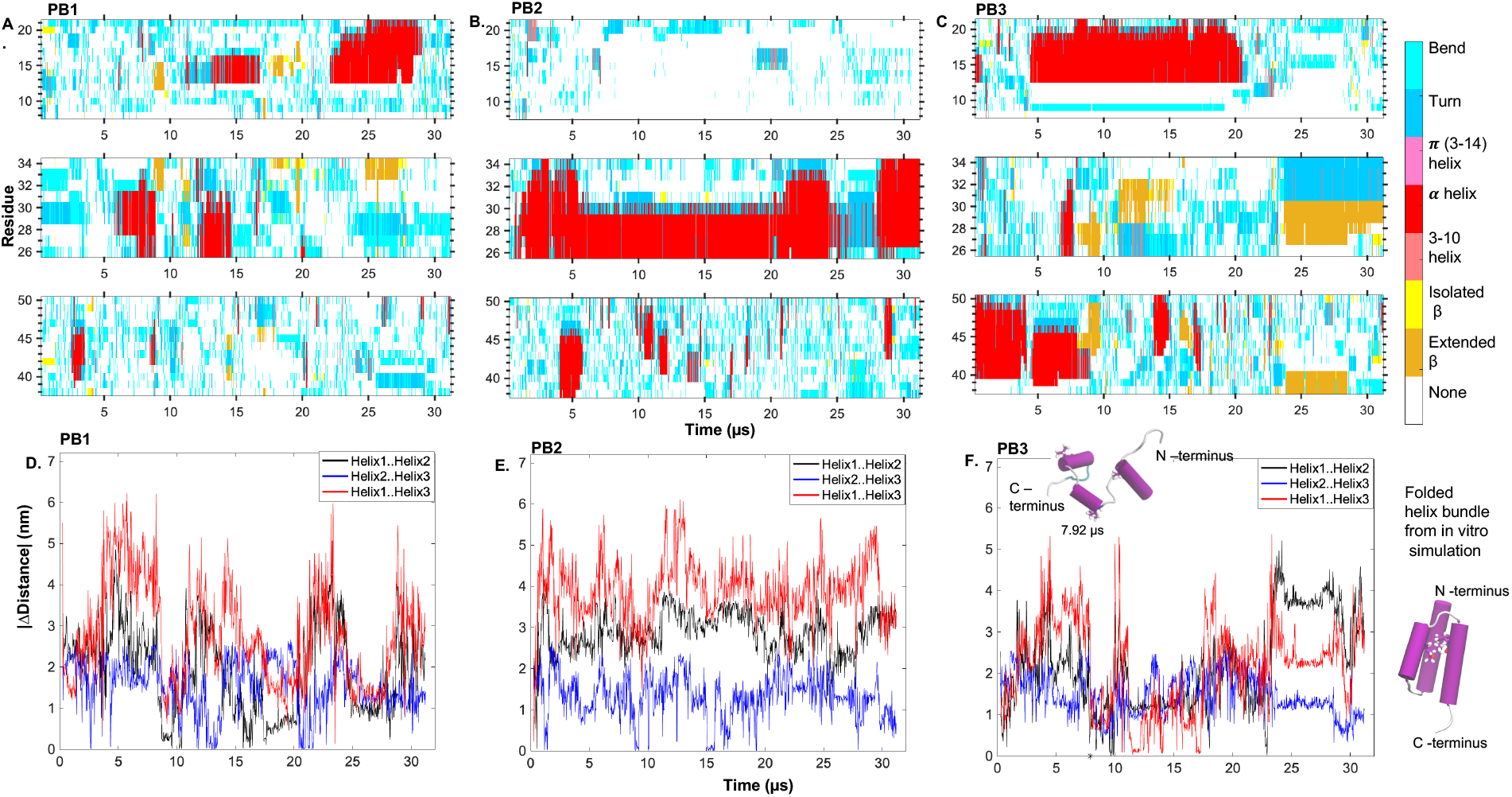
Secondary and tertiary structure formation in PB copies across the cell cytoplasm simulation. Formation of Helix1 (Lys8 to Ala21) – 1^st^ row, Helix2 (Asp26 to Lys34) – 2^nd^ row, and Helix3 (Val38 to Lys50) – 3^rd^ row in **(A)** PB1; **(B)** PB2; and (**C)** PB3. Complete secondary structure evolution is shown in Fig. S1. Absolute relative distances between pairs of helix strands, with respect to folded PB, in **(D)** PB1; **(E)** PB2; and **(F)** PB3. Ala14 on Helix1, Ile32 on Helix2, and Val41 on Helix3 were chosen as probes (represented as licorices in the cartoon figures above) to calculate distances between helices (center-of-mass distance between the probe sidechains). The marker * in panel F represents the 7.92 μs time point where PB3 adopted native-like secondary structure, but not the native topology (cartoon inset). The folded PB structure on the right was obtained following minimization and 20 ns MD simulation in an *in vitro* water box balanced with counter ions. Secondary structure elements were calculated using the DSSP algorithm^18^.

**Fig. 4.**
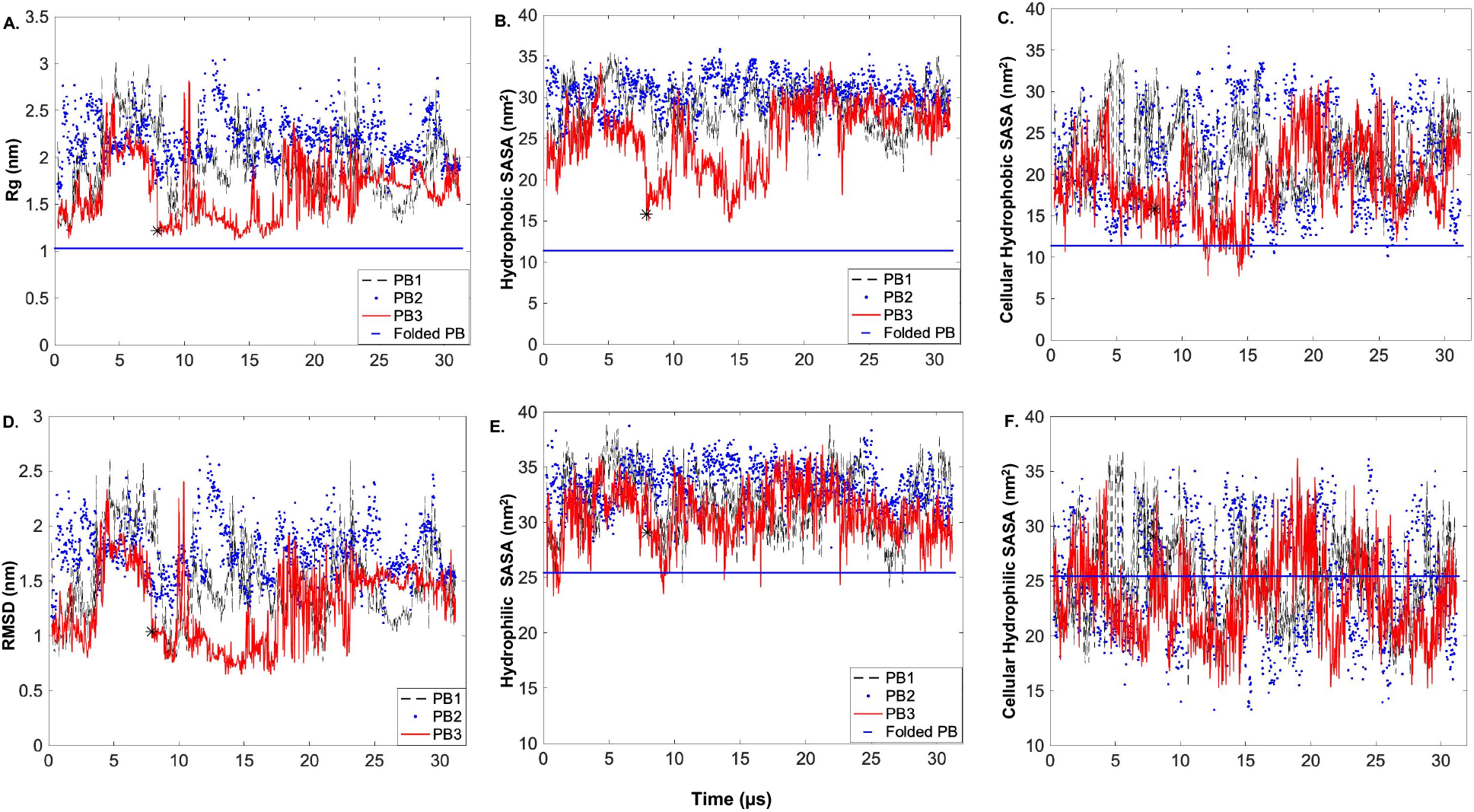
Characterizing protein dynamics of PB1/2/3 across the cytoplasm simulation. **(A)** R_g_, **(B)** hydrophobic SASA, **(C)** cellular hydrophobic SASA, **(D)** C-α RMSD, **(E)** hydrophilic SASA, and **(F)** cellular hydrophilic SASA of PB copies. Cellular SASAs for PBs in **(C)** and **(F)** were calculated by considering shielding effects provided by the surrounding solutes in the cytoplasm model. SASAs in **(B)** and **(C)** were calculated for individual PBs without considering the neighboring cellular solutes. Reference folded PB structure was obtained as described in Fig. 3. The data point * in each panel represents the almost folding event of PB3 towards native state at 7.92 μs.

In fact, PB3 quickly forms two anti-parallel β strands, consisting of Phe27-Phe29 and Ala43-Val45, while undergoing sticking interactions with cyclophilin G (CycG) protein at the partially unfolded region of Helix 1 N-terminal (Fig. 1B). Ala43, Leu 44, and Tyr28 are among the frequently interacting residues with CycG (Fig.5A, Table S3). Stabilizing interactions between the β barrel of CycG and the β strand elements in PBs were likely occurring across the simulations via both long-range attractive and cross-strand side-chain interactions (Figs. 2B, 5A)^15,16^. When helix propensity was scored^17^ for the three helix sequences of PB (Table S1), Helix1 in PB had the highest average helix propensity. Consequently, the maximum lifetime of β strands in Helix1 is much shorter across the different PB copies during the cytoplasm simulation. However, about 80% of Helix1 remained disordered for ~10 μs in PB2.

**Fig. 5.**
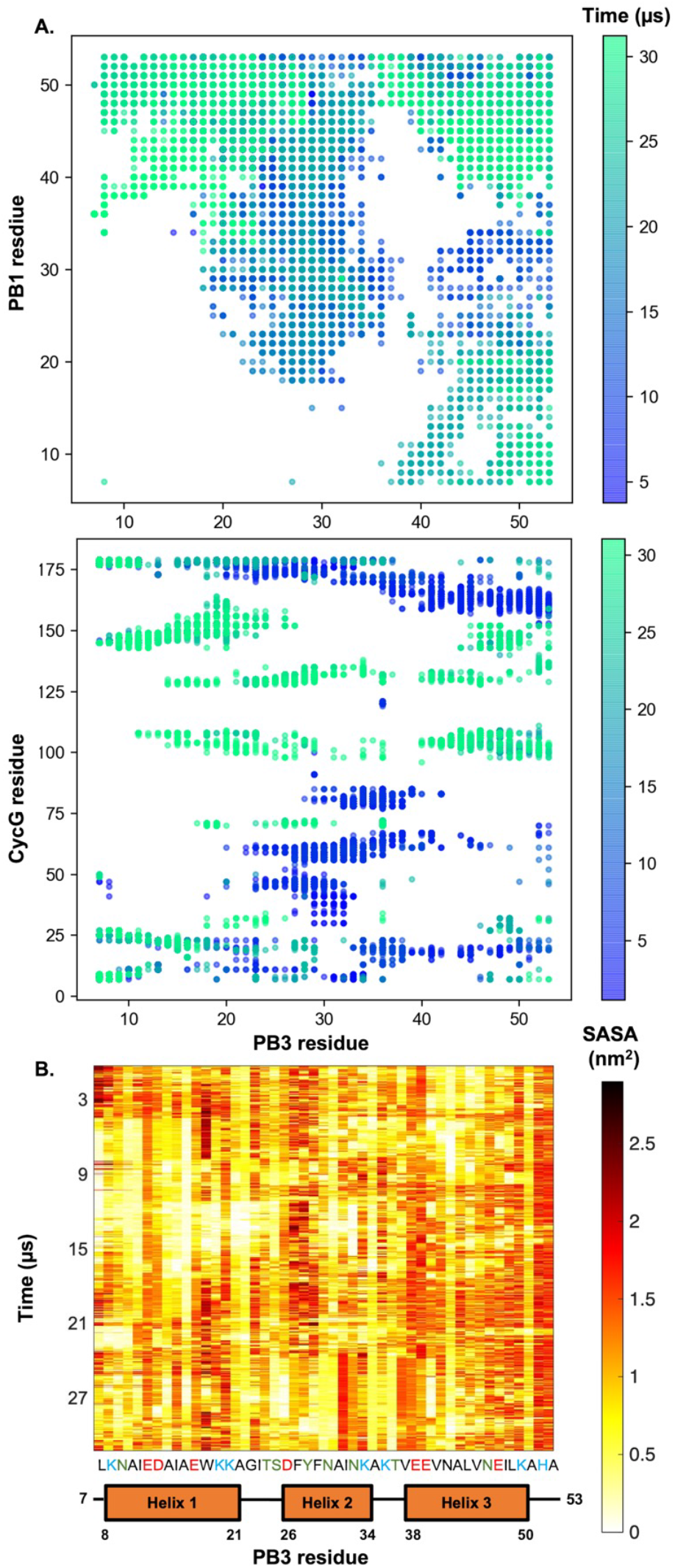
Contact maps across the cytoplasm simulation. **(A)** Time evolvement of residue specific sticking (involving CycG) and aggregating interactions (involving PB1) for PB3. These two PB3 interacting protein partners had the largest total contact surface area with PB3 during simulations. **(B)** Residue specific cellular SASA for PB3.

Seeding of β strands in PB also occurred when PB1 and PB3 interacted, forming a homodimeric proto-aggregate (Fig. 2). In panel 2 of Fig. 2A, we observed an anti-parallel β sheet formed comprised of Val41-Leu44 on PB1 interfacing with Gly22-Ser25 on PB3, which is also interfacing with Phe29-Lys34 on another PB3 strand. In the following frames, anti-parallel β strands were seen forming between Lys50-Ala53 on PB1 and Val41-Leu44 on PB3. PB1 and PB3 were also seen interacting through β strands while interacting with CycG in Fig 2B. Thus, protein sticking can lead to a chain of events whereby protein copies stick to other macromolecules in the cell, acquire β strand structure, and then aggregate with one another. A past study involving dynamic light scattering showed PB protein to exist in a mixture of homo-oligomeric states in solution, with the dominant species being a homo-tetramer^19^. Given our present observation of initial aggregating events between PB proteins, these homo-tetramers could accelerate aggregate formation in-cell.

In contrast, the binding of PB and human serum albumin (HSA) does not involve any fold changes in PB. Some of the PB residues that interface with HSA – including Ser25, Tyr28, Ala31, and Leu44 (Tables S2, S3)^20^ – have been observed to be among the top residues involved in sticking (with CycG) or aggregating (between PB copies) interactions described in the present study.

Within an *in vitro* environment, protein folding kinetics and thermodynamics are primarily driven by the hydrophobic effect, when non-polar residues are buried away from solvent^21^. Within the crowded cell, the attractive sticking interactions (electrostatic and hydrophobic) between macromolecules compete with hydrophobic burial and lead to a more rugged protein folding energy landscape. When we calculated the hydrophobic solvent accessible surface area (SASA) for the PB copies across the simulation trajectories by excluding the surrounding cellular solutes, the SASAs were at least 5 nm^2^ larger relative to the folded PB in an *in vitro* environment (Fig. 4B).

However, when the PB hydrophobic SASA was measured by considering the shielding interactions provided by the surrounding solutes, we observe the resulting ‘cellular hydrophobic SASAs’ for all the PB copies shifted toward the folded PB hydrophobic SASA (Fig. 4C) value in vitro. When hydrophilic SASAs were measured for the PBs (Figs. 4E,4F) in a similar manner, the cellular hydrophilic SASAs of the PBs fluctuated within the range of 10 nm^2^ above and below the folded PB hydrophilic SASA. Hydration of the hydrophilic surface of a protein provides stabilizing interactions for the protein, and can guide protein folding by mediating interactions between polar or charged residues^22^. Here, when cellular solute interactions are considered, the effect of hydration on protein folding is seen to compete with the sticking interactions between PBs and the surrounding biomolecules. In Fig. 5B, charged residues such as Lys8 and Lys20 in PB3, for example, are seen to have low solvent exposure for extended periods, and are also among the top sticky residues with CycG (Fig 5A, Table S3).

PB protein has the GA domain structure, known to be able to undergo fold switching when mutated^23^. A theoretical study hypothesized a fold switch can be triggered just by fatty acid binding around the albumin binding site in GA domain^24^. The sticking interactions between cellular macromolecules and PB copies are driving a wide range of α and β structural elements being explored in our study. Such structural plasticity could facilitate fold switching *in vivo,* and should be investigated further.

## Acknowledgements

Anton 2 computer time was provided by the Pittsburgh Supercomputing Center (PSC) through Grant R01GM116961 from the National Institutes of Health. T.V.P. acknowledges support from the NIH grant R01-GM141298 and the Department of Chemistry, University of Illinois at Urbana-Champaign. P.S.R., M.R., M.B., and M.G. were supported by the NSF grant MCB 2205665.

The authors declare no competing financial interests.

## Supplementary information

### 1. Methods

Computational methods of the cytoplasm model setup and simulation has been described in our previous study^8^. VMD, CPPTRAJ, and MDtraj packages were used for data analysis. Production run of the simulation was done on PSC Anton 2 computing resource.

### 2. PB sequence used in our study

Amino acids are colored black for hydrophobic, green for polar, blue for positively charged, and red for negatively charged. The initial 6 residues at the N-terminal in the protein B sequence from *Peptostreptococcus magnus* is deleted here.

**Fig. S1.**
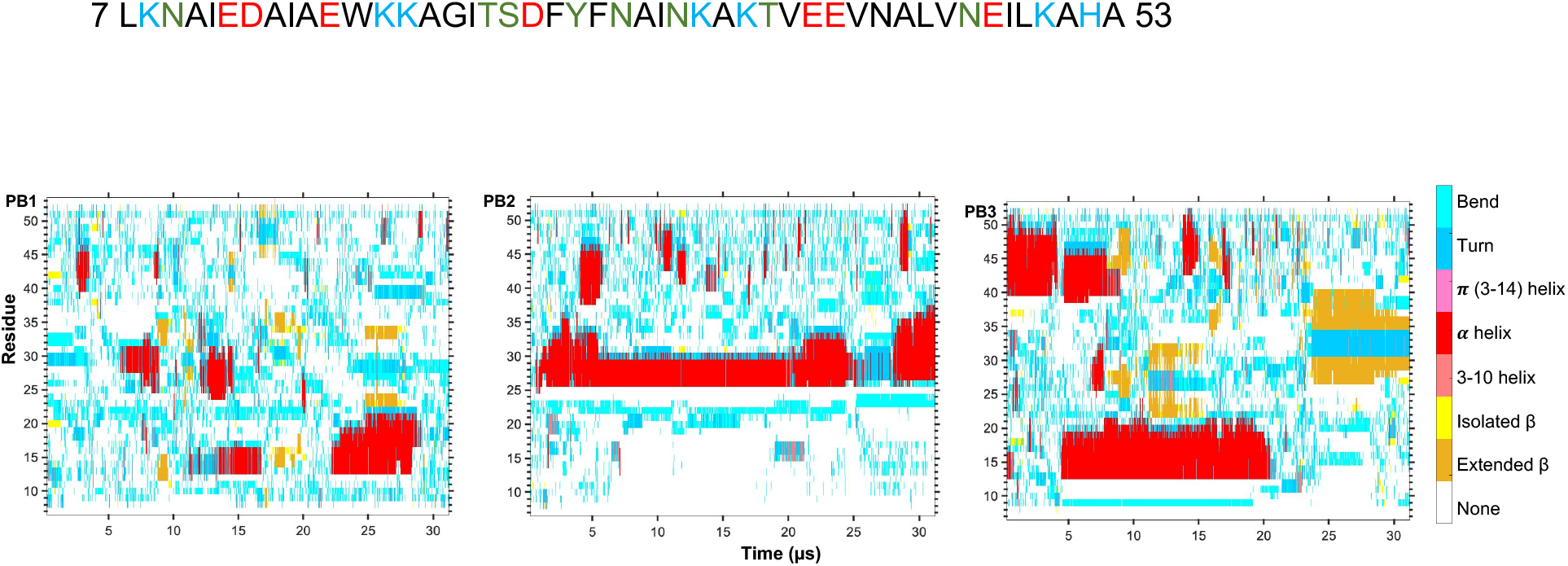
Secondary structure elements in PB copies across cytoplasm simulation. Calculations done using DSSP^11^ algorithms.

**Fig. S2.**
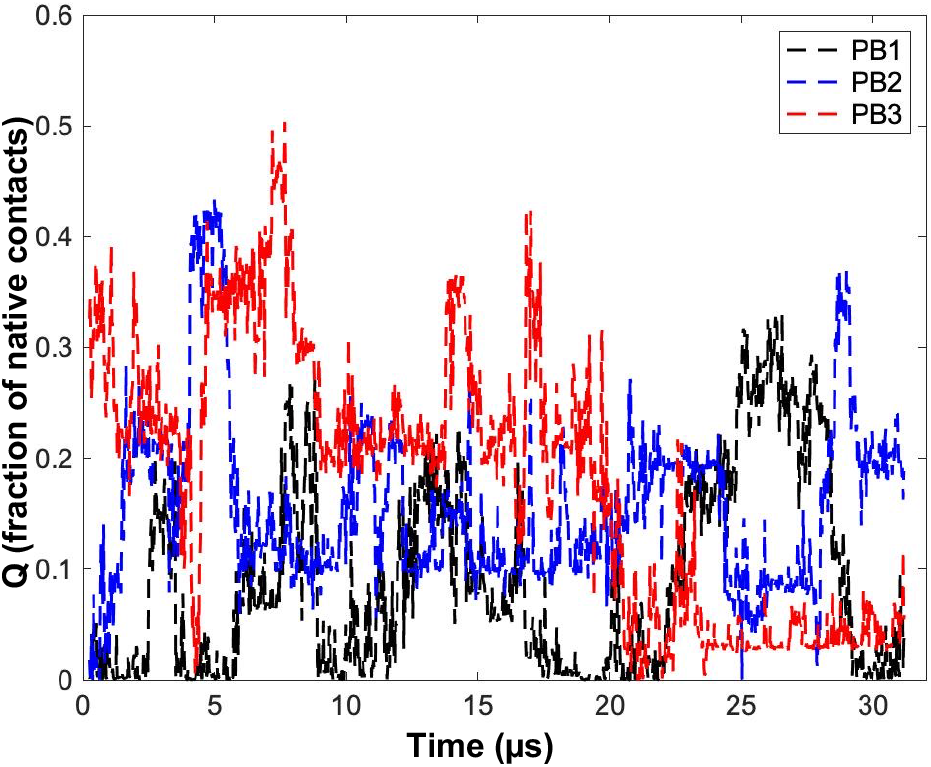
Fraction of native contacts across PB copies.

**Table S1.** Helix Propensity^14^ Table for PB

| Helix | Sequence | Average helix propensity score | Net charge, pH 7.4 |
| --- | --- | --- | --- |
| H1 | KNAIEDAIAEWKKA | 0.302 | -0.851 |
| **H2** | DFYFNAINK | 0.47 | -0.338 |
| **H3** | VEEVNALVNEILK | 0.415 | -2.475 |

**Table S2.** Most frequent (top 50) residue specific aggregating interactions between PB1 and PB3 during cytoplasm simulation.

| **PB1/PB3** | **Contact frequency** | **PB3 Helix (H)** |
| --- | --- | --- |
| LYS36/LYS8 | 0.1626646 | H1 |
| ALA43/LEU44 | 0.11541441 | H3 |
| LEU44/LEU44 | 0.10534469 | H3 |
| ILE48/ILE11 | 0.09837335 | H1 |
| ASN42/LEU44 | 0.09372579 | H3 |
| PHE29/ALA31 | 0.09062742 | H2 |
| LEU49/TRP18 | 0.08907823 | H1 |
| LYS50/ASP13 | 0.08365608 | H1 |
| PHE29/ASN30 | 0.08210689 | H2 |
| PHE29/ILE32 | 0.0813323 | H2 |
| ASN46/ALA21 | 0.0813323 | H1 |
| ASN30/PHE29 | 0.07978311 | H2 |
| ASN46/GLY22 | 0.07823393 |  |
| ASN30/ASN30 | 0.07745933 | H2 |
| LYS36/LEU7 | 0.07668474 |  |
| LEU49/ALA21 | 0.07591015 | H1 |
| ILE48/GLY22 | 0.07126259 |  |
| ILE48/ASN9 | 0.0697134 | H1 |
| ILE48/ILE23 | 0.06893881 |  |
| VAL45/ALA21 | 0.06738962 | H1 |
| PHE29/PHE29 | 0.06661503 | H2 |
| VAL45/GLY22 | 0.06506584 |  |
| GLU47/GLY22 | 0.06506584 |  |
| LEU49/GLY22 | 0.06506584 |  |
| LEU49/ILE23 | 0.06506584 |  |
| ALA51/ALA14 | 0.06351665 | H1 |
| LYS50/ALA14 | 0.06351665 | H1 |
| ALA51/GLU12 | 0.06351665 | H1 |
| LEU49/ILE11 | 0.06274206 | H1 |
| LYS50/GLU17 | 0.06196747 | H1 |
| GLU47/ALA21 | 0.06196747 | H1 |
| ILE48/ALA21 | 0.06196747 | H1 |
| ILE32/PHE29 | 0.06119287 | H2 |
| LEU49/THR24 | 0.06119287 |  |
| ASP26/LYS34 | 0.05964369 | H2 |
| LYS50/GLU12 | 0.05886909 | H1 |
| LEU44/VAL45 | 0.05886909 | H3 |
| ASN30/ALA31 | 0.0580945 | H2 |
| ILE48/THR24 | 0.05731991 |  |
| VAL45/LEU44 | 0.05577072 | H3 |
| LEU44/ASN46 | 0.05577072 | H3 |
| ALA51/ASP13 | 0.05422153 | H1 |
| ASN46/LYS20 | 0.05344694 | H1 |
| ILE48/LYS8 | 0.05267235 | H1 |
| VAL45/GLU47 | 0.05034857 | H3 |
| LYS50/ILE11 | 0.05034857 | H1 |
| ALA43/VAL45 | 0.05034857 | H3 |
| TYR28/ILE32 | 0.04957397 | H2 |
| VAL41/PHE29 | 0.04879938 | H2 |

**Table S3.** Most frequent residue specific sticking interaction between CycG and PB3 during cytoplasm simulation.

| **CycG/PB3** | **Contact frequency** | **PB3 Helix (H)** |
| --- | --- | --- |
| ARG9/LEU7 | 0.65453137 |  |
| ARG9/LYS8 | 0.64368706 | H1 |
| LEU177/ASN9 | 0.56312936 | H1 |
| PRO179/LEU7 | 0.55925639 |  |
| ARG7/LEU7 | 0.54144074 |  |
| LEU177/LEU7 | 0.49186677 |  |
| LEU177/ILE11 | 0.48179706 | H1 |
| VAL25/ASN9 | 0.47947328 | H1 |
| ILE145/LYS8 | 0.47560031 | H1 |
| ILE178/LEU7 | 0.47560031 | H1 |
| SER146/ASN9 | 0.47017816 | H1 |
| VAL25/LYS8 | 0.45701007 | H1 |
| SER146/LYS8 | 0.45546088 | H1 |
| PRO8/LEU7 | 0.42912471 |  |
| LEU177/ALA10 | 0.41518203 | H1 |
| ARG23/ILE11 | 0.38032533 | H1 |
| ILE178/LYS19 | 0.37412858 | H1 |
| SER146/ALA10 | 0.37103021 | H1 |
| ILE178/LYS20 | 0.36948102 | H1 |
| ARG23/ALA10 | 0.36793184 | H1 |
| ARG7/LYS8 | 0.36560806 | H1 |
| ILE178/ILE11 | 0.36405887 | H1 |
| ILE178/ALA16 | 0.35941131 | H1 |
| PRO179/LYS19 | 0.35786212 | H1 |
| VAL25/LEU7 | 0.32300542 |  |
| VAL25/ALA10 | 0.30906274 | H1 |
| LEU177/LYS8 | 0.30906274 | H1 |
| ARG23/ASP13 | 0.28505035 | H1 |
| ARG23/GLU12 | 0.28505035 | H1 |
| PHE11/ALA10 | 0.22850503 | H1 |
| GLU27/LYS8 | 0.19674671 | H1 |
| ASN105/ALA43 | 0.16731216 | H3 |
| PRO130/ALA43 | 0.1626646 | H3 |
| PRO132/PHE29 | 0.16189001 | H2 |
| HSD104/LEU44 | 0.16111541 | H3 |
| ILE145/ASN9 | 0.15956623 | H1 |
| ASP71/LYS36 | 0.15879163 |  |
| ASN105/LEU44 | 0.15646785 | H3 |
| HSD104/ALA43 | 0.15646785 | H3 |
| LYS129/ASP26 | 0.15414407 | H2 |
| ASN105/ASN42 | 0.15336948 | H3 |
| SER146/ILE11 | 0.15182029 | H1 |
| ARG23/ASN9 | 0.1510457 | H1 |
| PRO132/TYR28 | 0.14949651 | H2 |
| ARG9/ASN9 | 0.14794733 | H1 |
| ASN105/VAL41 | 0.14252517 | H1 |
| PRO130/VAL41 | 0.1394268 | H1 |
| LYS129/SER25 | 0.13323005 |  |
| PRO132/VAL41 | 0.13245546 | H3 |
| HSD104/ASN42 | 0.13090627 | H1 |

